# Mitochondrial diabetes in mice expressing a dominant-negative allele of nuclear respiratory factor-1 (*Nrf1*) in pancreatic β-cells

**DOI:** 10.1101/2023.01.22.524153

**Authors:** Fionnuala Morrish, Helene Gingras, Joanna Noonan, Li Huang, Ian R. Sweet, Iok Teng Kuok, Sue E. Knoblaugh, David M. Hockenbery

## Abstract

Genetic polymorphisms in nuclear respiratory factor-1 (*NRF1*), a key transcriptional regulator of nuclear-encoded mitochondrial proteins, have been linked to diabetes. Homozygous deletion of *Nrf1* is embryonic lethal in mice. Our goal was to generate mice with β-cell-specific reduction in NRF1 function to investigate the relationship between NRF1 and diabetes. We report the generation of mice expressing a dominant-negative allele of *Nrf1* (DNNRF1) in pancreatic β-cells. Heterozygous transgenic mice had high fed blood glucose levels detected at 3 wks of age, which persisted through adulthood. Plasma insulin levels in DNNRF1 transgenic mice were reduced, while insulin sensitivity remained intact in young animals. Islet size was reduced with increased numbers of apoptotic cells, and insulin content in islets by immunohistochemistry was low. Glucose-stimulated insulin secretion in isolated islets was reduced in DNNRF1-mice, but partially rescued by KCl, suggesting that decreased mitochondrial function contributed to the insulin secretory defect. Electron micrographs demonstrated abnormal mitochondrial morphology in β- cells. Expression of NRF1 target genes *Tfam*, *T@1m* and *T@2m*, and islet cytochrome c oxidase and succinate dehydrogenase activities were reduced in DNNRF1-mice. Rescue of mitochondrial function with low level activation of transgenic c-Myc in β-cells was sufficient to restore β-cell mass and prevent diabetes. This study demonstrates that reduced NRF1 function can lead to loss of β-cell function and establishes a model to study the interplay between regulators of bi-genomic gene transcription in diabetes.

## INTRODUCTION

In the pancreatic β-cell, mitochondrial function is required for the coupling of glucose metabolism to insulin secretion (Maechler et al., 2006; Jensen et al., 2008; Matschinsky and Collins, 1997; Wiederkehr and Wollheim, 2012). Extensive evidence shows that reduction of mitochondrial activity in islets and insulin-secreting cell lines by a variety of insults, including inhibition of cytoplasmic NADH shuttles or cataplerotic/anaplerotic functions, and lipotoxicity, inhibits glucose-stimulated insulin secretion (GSIS) (Maedler et al., 2001; Antinozzi et al., 2002; Rubi et al., 2004; Noda et al., 2002; Campbell and Newgard, 2021). The effects of mitochondrial dysfunction reflect the role of mitochondria in β−cell ATP generation, as increased energy charge (ATP/ADP) is a key factor in closing KATP channels that leads to elevated flux of calcium through L-type calcium channels and stimulation of trafficking and exocytosis of insulin granules. Decreased ATP production therefore will result in diminished insulin secretion.

Pancreatic β-cell dysfunction is observed with mutations in mitochondrial DNA in some diabetes patients (Kobayashi et al, 1997; Katagiri et al., 1994; Bensch et al., 2009; Karaa and Goldstein, 2015). Transcriptional regulators of mitochondrial biogenesis include peroxisome proliferator-activated receptor gamma coactivator 1 alpha, beta and PGC-1-related coactivator (PGC-1α, PGC-1β, PRC), nuclear respiratory factors 1 and 2 (NRF1, NRF2), and estrogen-related receptor alpha (ERRα) (Handschin and Spiegelman, 2006; Scarpulla, 2008). Alterations in the expression of these transcription factors may also occur in diabetic islets. For example, PGC-1α expression is reduced in islets from Type 2 diabetes (T2D) patients (Ling et al., 2008).

An important downstream target of these mitochondrial regulators is the transcription factor *TFAM*, a nuclear-encoded gene that regulates the transcription and replication of mitochondrial DNA (Kang et al., 2007). Targeted knockout of *Tfam* in β-cells results in impaired insulin secretion and β cell loss (Silva et al., 2000). Diabetes in this model was transient as mice older than 20 wks showed normalization of fasting blood glucose due to incomplete knockout in β-cells.

Many of the effects of PGC-1α and related co-activators are mediated through activation of the transcription factor NRF1 (Scarpulla, 2008). NRF1 regulates genes required for mitochondrial biogenesis, including *TFAM*, *TFB1M* and *TFB2M* transcription factors, and subunits of the respiratory complexes of the electron transport chain, including all 10 nuclear-encoded genes for cytochrome c oxidase (Gleyzer et al., 2005; Dhar et al., 2009). There are several reports linking genetic polymorphisms in *NRF1* with type 2 diabetes (Cho et al., 2005; Gaulton et al., 2008; Liu et al., 2008). However, the role of NRF1 in β-cell function is unknown, as *Nrf1* knockout in mice is embryonic lethal between day E3.5 and E6.5 with decreased cell proliferation and mtDNA content (Huo and Scarpulla, 2001). To address the role of Nrf1 in regulating mitochondrial function in pancreatic β-cells, we have generated transgenic mice that selectively express a dominant-negative allele of *Nrf1* (DNNRF1) in pancreatic β-cells under the control of the rat *Ins2* promoter. *pInsDNNRF-1ER^TAM^* transgenic mice develop diabetes at an early age that persists over the lifetime of the mouse and is associated with weight loss, decreased serum insulin and β-cell loss due to apoptosis. This model provides further genetic evidence of a critical role for mitochondria in normal glucose-stimulated insulin secretion and suggests a role for NRF1 as a key regulator of mitochondrial biogenesis in β-cells.

## RESULTS

### Generation of transgenic mice expressing dominant-negative Nrf1

Using a previously published human dominant-negative *NRF1* construct with a deleted transactivation domain (DNNRF1), we generated the plasmid *pIns-DNNRF1ER^TAM^* containing an in-frame fusion to the estrogen receptor under control of the rat *Ins2* promoter (Figure 1A) (Morrish et al., 2003; Wu et al., 1999; Gonen and Assaraf, 2010; Tokusumi et al., 2004; Kherrouche et al., 2004). Founder mice were screened by PCR amplification of a region spanning the 3’ end of DNNRF1 and the 5’ end of the fused estrogen receptor. A total of 3 positive founders were generated and two successfully established founder lines (#2 and #3). Southern analysis was used to demonstrate transgene integration and copy number (Figure 1B). Copy numbers were estimated to be 10 and 1-2 by Southern blot for founders #2 and #3, respectively (data shown for founder #2 in Figure 1B). The transgene was transmitted to 50% of progeny. Heterozygous pups were normal in size compared to their wild-type littermates.

**Figure 1.**
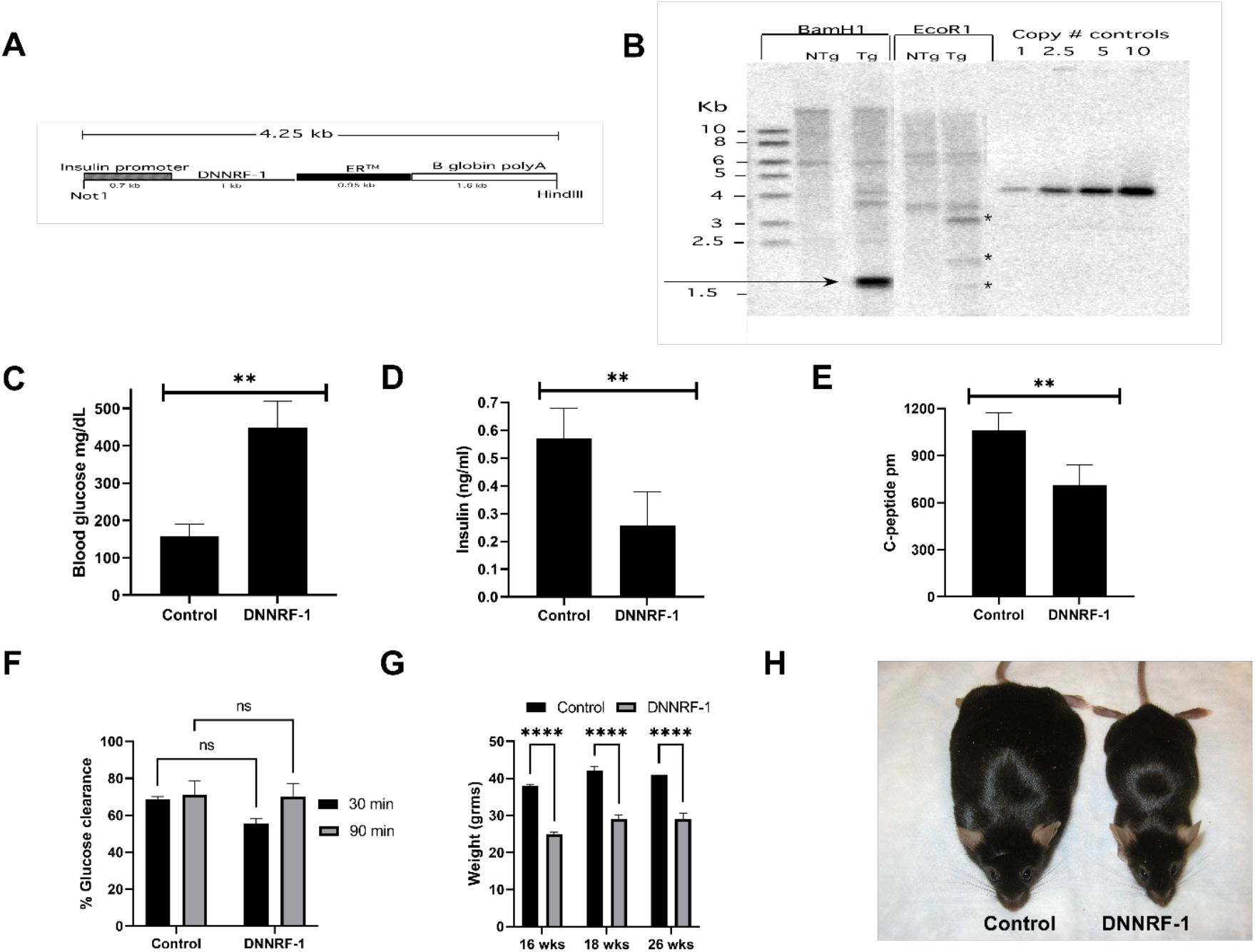
Transgene detection and functional analysis of physiological effects. (A) pIns-*DNNRF-1ER^TAM^* transgene cassette. The 4.25 kb transgene cassette containing the Ins-1 promoter, *DNNRF-1ER^TAM^* fusion gene and β globin polyA sequences. (B) Southern blot analysis of copy number. Includes non-transgenic (NTg) and transgenic (Tg). (C) Fasting blood glucose levels for control (non-transgenic) and transgenic pIns-*DNNRF-1ER^TAM^* (DNNRF-1) mice at 8 wk of age (n=9). (D) Fasting serum insulin concentrations for control and transgenic pIns-*DNNRF-1ER^TAM^* (DNNRF-1) mice at 8 wk of age (n=9). (E) Serum C-peptide concentrations for control and transgenic pIns-*DNNRF-1ER^TAM^* (DNNRF-1) mice at 8 wk of age (n=9). (F) Glucose response of mice at 30 minutes and 90 minutes after administration of 1 mU/g insulin i.p, expressed as percent of baseline, at 6 months of age (n=4). (G) Weight of control and transgenic mice at 10, 16 and 26 weeks (n=5). (H) Photograph of control and DNNRF-1 mice at 26 weeks.

### *pIns-DNNRF1ER^TAM^* transgenic mice are diabetic

Fasting hyperglycemia (459.2+/-40.0 mg/dl; 497.4+/-29.8 mg/dl) and glucosuria (data not shown) were observed in both founder lines of transgenic *pIns-DNNRF1ER^TAM^* mice at 6-8 wks of age without tamoxifen treatment (Figure 1C). Analysis of fed glucose levels over time detected hyperglycemia as early as 3 wks of age (Supplemental Figure 1A). This suggested that leaky activation of the DNNRF1ER^TAM^ protein was sufficient to induce diabetes and all further studies were conducted in the founder line #2 in the absence of tamoxifen. Fasting blood glucose data are shown for males in Figure 1C; females also had a diabetic phenotype (mean 428+/-32 mg/dL) with a slightly lower mean fasting glucose. Fasting serum insulin concentrations were significantly less than control levels in 8 wk old transgenic mice (control 0.57+/- 0.108 ng/ ml, DNNRF1 0.257 +/- 0.122 ng/ml, p=0.013) (Figure 1D). Significant reductions were also observed in serum C-peptide concentrations (control 1060 +/- 112 pM, DNNRF1 709 +/- 132 pM, p=0.012) (Figure 1E). Treatment with insulin demonstrated normal insulin sensitivity at 2 months of age (Figure 1F), but insulin resistance was present at 9 months (data not shown). At 26 wk of age, transgenic mice showed persistently elevated fasting blood glucose (control 163+/- 9.8 mg/ dL, DNNRF1 > 600 mg/dL) and a significant reduction in body weight (control 41+/- 1.5 gm, DNNRF1, 29+/- 1.7 gm, p=0.013) (Figure 1G, H).

### Pancreatic β-cell depletion in Ins-*DNNRF-1ER^TAM^* mice

We analyzed histological sections from <u>pancreata</u> of *pIns-DNNRF1ER^TAM^* mice presenting with hyperglycemia at two months of age. H&E-stained sections showed small, deformed islets (Figure 2A, B). To determine if the alterations in islet morphology in *pIns-DNNRF1ER^TAM^* mice were associated with increased apoptotic cell death, we immunostained paraffin-fixed pancreatic sections for cleaved caspase-3. Numerous caspase-3-positive cells were present within islets from *pIns-DNNRF1ER^TAM^* mice, while caspase-3 staining in wild-type littermates was confined to a few peripheral islet cells (Figure 2C, D), as previously reported (Ladiges et al., 2005).

**Figure 2.**
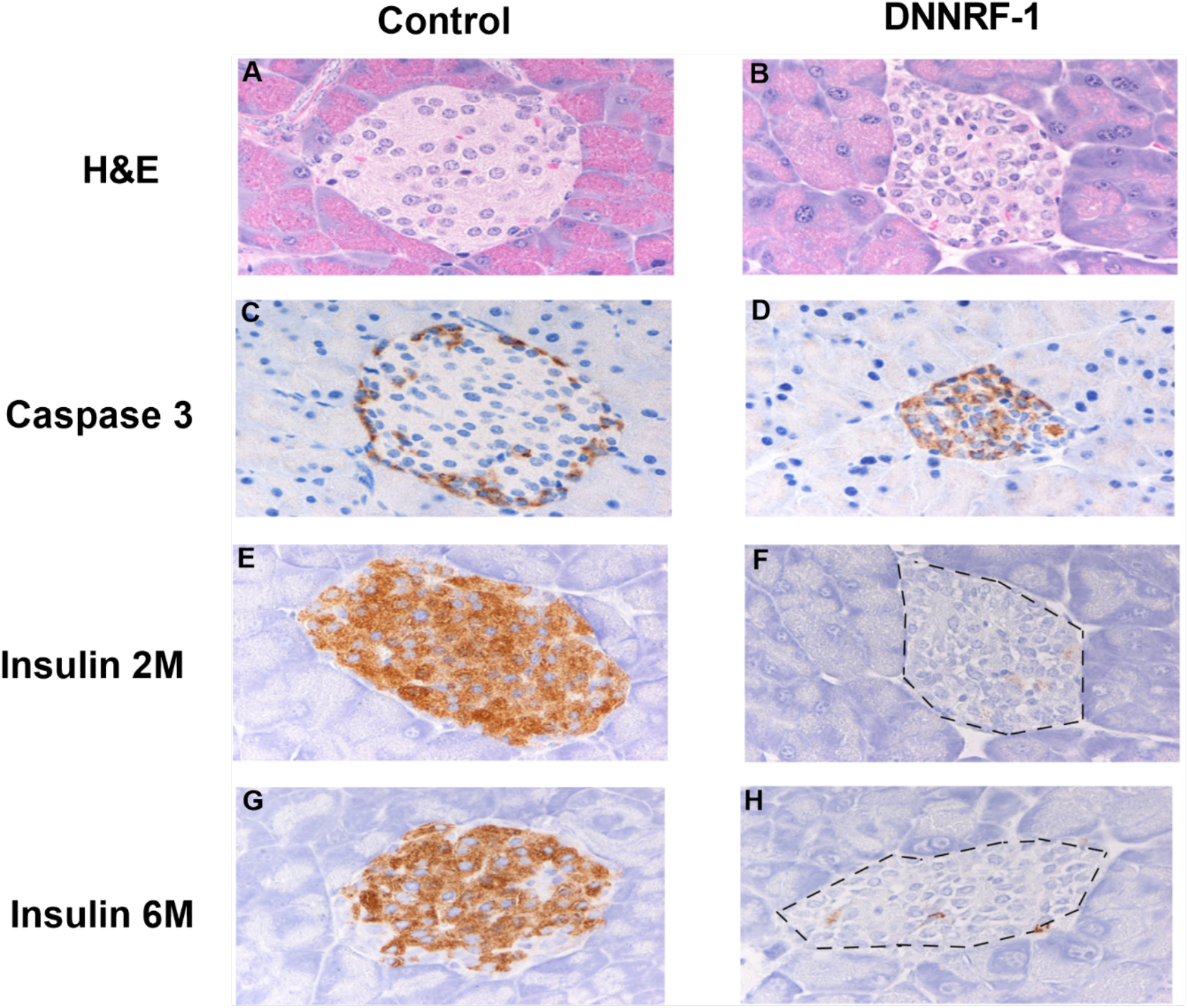
**Histology and immunohistochemistry studies of pancreatic tissue.**(A, B) H&E staining of pancreatic tissue of control (A) and pIns-*DNNRF-1ER^TAM^* mice (B). (C, D) Activated Caspase 3 immunostaining of pancreatic islets of control (C) and pIns-*DNNRF-1ER^TAM^* mice (D). (E, F, G, H) Insulin immunostaining of pancreatic islets for control (E) and pIns- *DNNRF-1ER^TAM^* mice (F) at 2 and 6 m of age (G, H). Photomicrographs are representative of n=4 or more mice from each group. All images were taken at 40 X magnification.

Staining of paraffin sections with an anti-insulin antibody showed sparse, scattered staining in *pIns-DNNRF1ER^TAM^* mice at two months of age (Figure 2E, F) compared to islets in wild-type littermates. The loss of insulin staining persisted in mice at 6 months of age (Figure 2G, H).

Quantitation of islet morphology demonstrated that average islet area was decreased (Supplemental Figure 1B) and individual islet cells were crowded, with increased nuclei per unit islet area (Supplemental Figure 1D). These pathological changes were limited to the islets, with no histological changes in pancreatic acini. Quantitative differences in apoptotic cells and insulin staining between wild-type and transgenic islets were statistically significant (Supplemental Figure 1C, E).

Staining with anti-glucagon antibody demonstrated a predominance of α-cells in islets from DNNRF1 transgenic mice (Supplemental Figure 2).

### DNNRF1 and target genes

We performed qRT-PCR to determine if DNNRF1ER^TAM^ altered the expression of *Tfam*, *T@1m*, and *T@2m*, NRF1 target genes involved in regulation of mitochondrial transcription. Expression of the three genes was significantly lower in DNNRF1 transgenics (>80% reduction; Supplemental Table 1). The *Ins2* promoter used in this study has been shown to result in transgene expression in the hypothalamus and at low levels, in kidney and duodenum, in addition to pancreatic β−cells (Wicksteed et al., 2010). Hypothalamic expression of *Tfam*, *T@1m* and *T@2m* was not significantly different between wild type and DNNRF1-mice (Supplemental Table 1). A non-mitochondrial target of NRF1 transactivation normally expressed in pancreatic β-cells, *Vsnl1*, was reduced in expression in DNNRF1 islets by immunohistochemistry (Supplemental Figure 3) (Fu et al., 2009; Dai et al., 2006).

### Mitochondrial function and morphology in *pIns-DNNRF1ER^TAM^* islets

To determine if the reduced insulin staining and altered morphology of *pIns- DNNRF1ER^TAM^* islets were linked to diminished NRF1-dependent respiratory chain function, we examined cytochrome c oxidase (COX) and succinate dehydrogenase (SDH) activities in sections using histochemical assays. NRF1 regulates the transcription of all 10 COX nuclear genes, and all 4 SDH subunits, as well as the mitochondrial transcription factor TFAM (Gleyzer et al., 2005; Dhar et al., 2008; Piantadosi and Suliman, 2008; Au and Scheffler, 1998). We observed a reduction in COX and SDH activities in islets of *pIns-DNNRF1ER^TAM^* mice at two months of age (Figure 3A-D).

**Figure 3.**
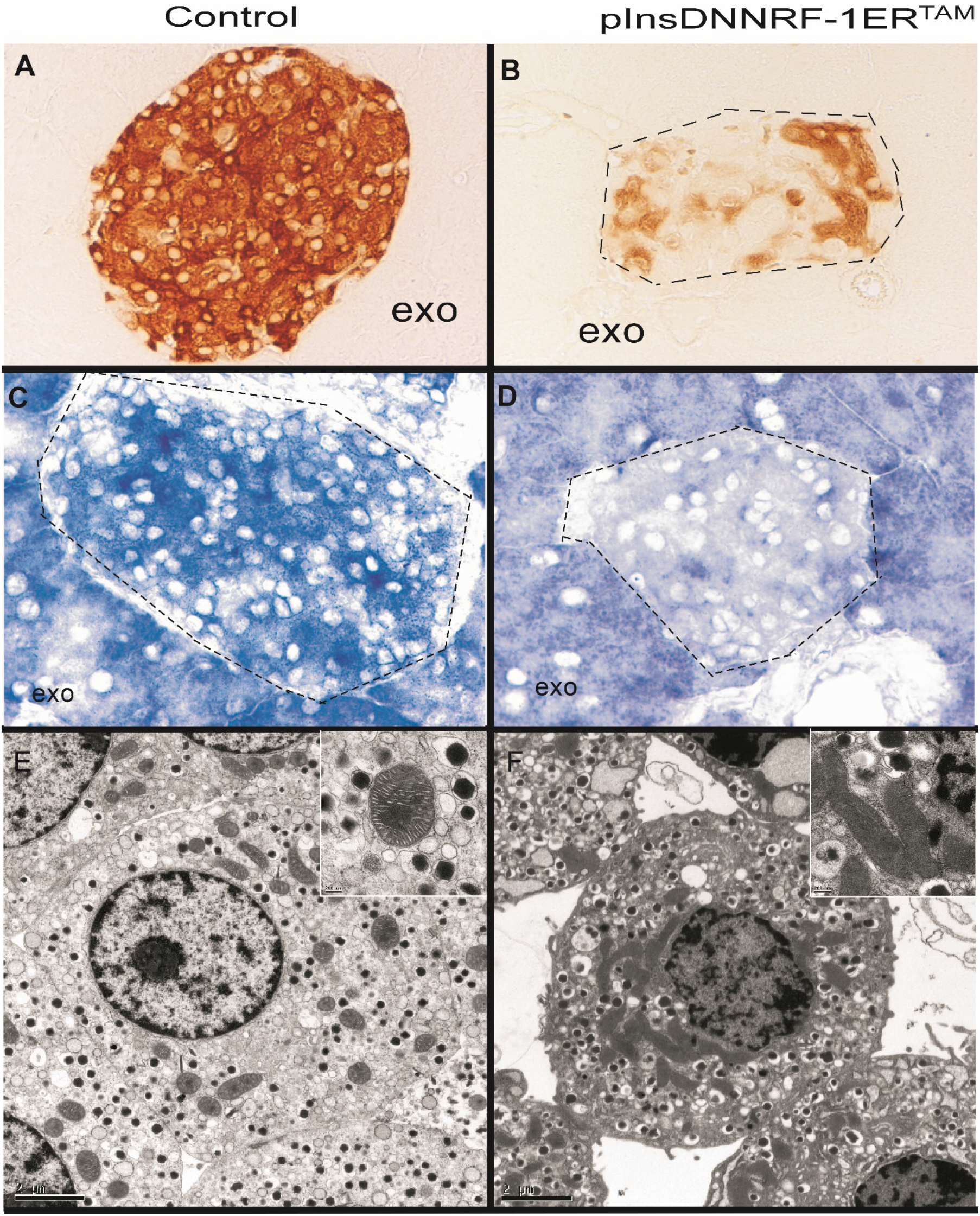
Cytochrome oxidase, succinate dehydrogenase histochemistry and electron microscopy of pancreatic islets. (A, B) Cytochrome c oxidase histochemical staining of pancreatic islets from control (A) and pIns- *DNNRF-1ER^TAM^* mice. (C, D) Succinate dehydrogenase histochemical staining for control (C) and pIns-*DNNRF-1ER^TAM^* mice (D). Photomicrographs of COX and SDH staining taken at 40 X magnification and representative of n=5 mice. Islets are outlined by dotted line and exocrine tissue labeled Exo. (E, F) Electron micrographs of pancreatic β cell from control (E) and pIns- *DNNRF-1ER^TAM^*. The bar in the electron micrographs is equivalent to 2 μm in the main panel and 200 nm in the insert.

In electron micrographs of isolated islets, cells were identified that exhibited the characteristic appearance of insulin secretory granules, with a central dense core and a surrounding halo (Figure 3E, F). In *pIns-DNNRF1ER^TAM^* mice, β-cells contained numerous elongated mitochondria with diffusely electron-dense matrix material obscuring the cristae (Figure 3F and insert). This appearance suggests accumulation of insoluble protein, possibly linked to incomplete assembly of ETC protein complexes (Morrish et al., 2003; Margineantu et al., 2007). β-cells in islets from both wild type and *pIns-DNNRF1ER^TAM^* mice were similar in size (Supplemental Figure 4A). A trend towards increased mitochondrial number and area in *pIns- DNNRF1ER^TAM^* mice did not reach statistical significance (Supplemental Figure 4B, C). However, we did find a significant increase in the number of secretory granules per cell, possibly consistent with a secretory defect (Supplemental Figure 4D).

Perfusion analysis of oxygen consumption rate (OCR) and glucose-stimulated insulin secretion (GSIS) was performed on isolated islets. DNNRF1 islets had significantly lower OCR in the presence of 3 mM glucose and this difference further increased with the addition of 20 mM glucose (Figure 4A). The GSIS was also significantly reduced, to less than 0.04 ng/min/100 transgenic islets, and there was no significant response to higher glucose concentrations (Figure 4B).

**Figure 4.**
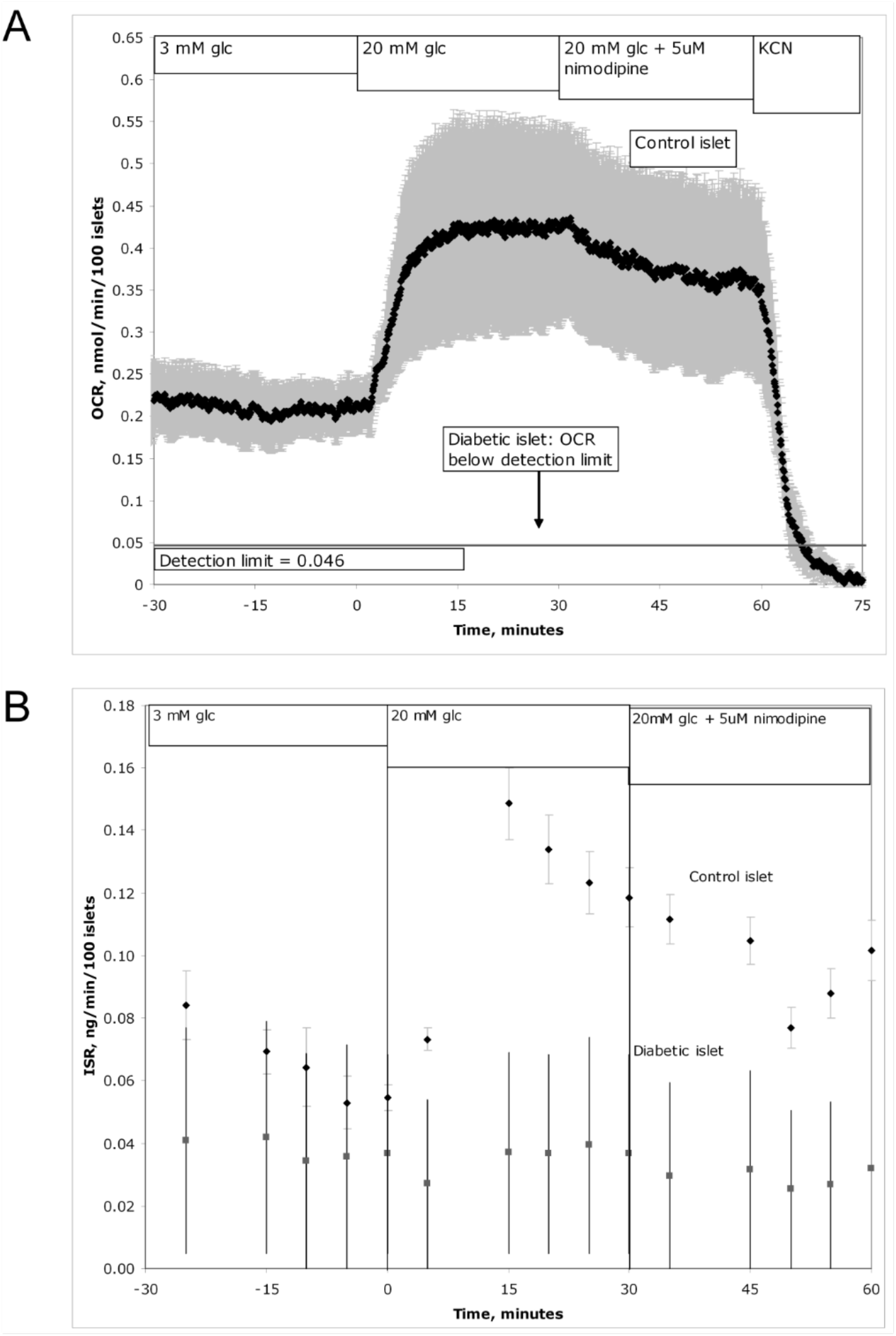
Glucose-stimulated oxygen consumption and insulin secretion from isolated islets. One hundred islets were perfused with 3 mM glucose for 30 min. The glucose in the perifusate was subsequently raised to 20 mM for 30 min, followed by sequential addition of 5 μM nimodipine and KCN. (A) Oxygen consumption rate (OCR) in isolated islets. The ANOVA p values were as follows; 3 mM glucose (p value<0.001), 20 mM glucose (p-value<0.001), 20 mM glucose+nimodipine (p-value>0.05). B) Insulin secretion rate (ISR) in isolated islets. Addition of the calcium channel blocker nimodipine inhibited Ca++-dependent OCR and KCN completely inhibited OCR in control islets. The ANOVA p-values were as follows; 3 mM glucose (p-value<0.05), 20 mM glucose (p-value<0.001), 20 mM glucose+nimodipine (p-value<0.001).

Static islet insulin assays confirmed a reduction in GSIS in DNNRF1 transgenics (Supplemental Figure 5). Insulin secretion was partially rescued in response to added KCl in DNNRF1 islets. These results are consistent with defective mitochondrial metabolism as a contributing mechanism of β-cell dysfunction in *pIns-DNNRF1ER^TAM^* mice. ATP levels (normalized to total protein) in isolated islets from *pIns-DNNRF1ER^TAM^* mice were less than 50% of control levels (Supplemental Figure 6).

### c-Myc transgene eliminates diabetes in *pIns-DNNRF1-ER^TAM^* mice

In previous work we showed that the c-Myc oncoprotein and NRF1 shared gene targets, due to common binding sites in promoters of a subset of nuclear-encoded mitochondrial genes (Morrish et al., 2003). Subsequent studies have confirmed c-Myc’s role in regulation of bi-genomic transcription and demonstrate that c-Myc, like NRF1, is involved in regulation of mitochondrial biogenesis (Li et al., 2005; Popay et al., 2021). c-Myc expression can trigger mitochondrial pathways of apoptosis under some conditions. Dual expression of c-Myc and DNNRF1 suppressed c-Myc-induced apoptosis in fibroblasts under nutrient-deprived conditions and improved mitochondrial function, suggesting that restoration of balanced mitochondrial gene expression and mitochondrial function opposes apoptosis (Morrish et al., 2003). To determine if c-Myc would likewise suppress DNNRF1-induced cytotoxicity in pancreatic β-cells, we generated *pIns-DNNRF1ER^TAM^ / pIns-c-MycER^TAM^* mice.

We found that the double transgenic mice had normal levels of fasting blood glucose, and normal responses to glucose challenge (Figure 5A, B). This result was observed without tamoxifen treatment, which is normally required for nuclear translocation of the c-MycER^TAM^ protein, suggesting leaky activation of c-MycER^TAM^ protein in β-cells (as shown in Pascal et al., 2008). The distribution of insulin staining in pancreatic islets was similar to controls (Supplemental Figure 7), confirmed as shown by quantitative image analysis (control insulin staining 92.6 +/- 1.0 %, *pIns-DNNRF1ER^TAM^* / *pIns-c-MycER^TAM^* 93.7 +/- 0.7%). Inducible activation of c-Myc in adult β cells produces rapid islet involution due to apoptosis, with almost complete ablation of β cells within 6-10 days (Pelengaris et al., 2002). Others have shown that, in the absence of tamoxifen, *pIns-c-MycER^TAM^* transgenic mice exhibit changes in β-cell proliferation and glucose-stimulated insulin secretion consistent with c-MycER^TAM^ activity (Pascal et al., 2008). This activity appears to be sufficient to rescue the negative effects of DNNRF-1 on glucose homeostasis and islet morphology.

**Figure 5.**
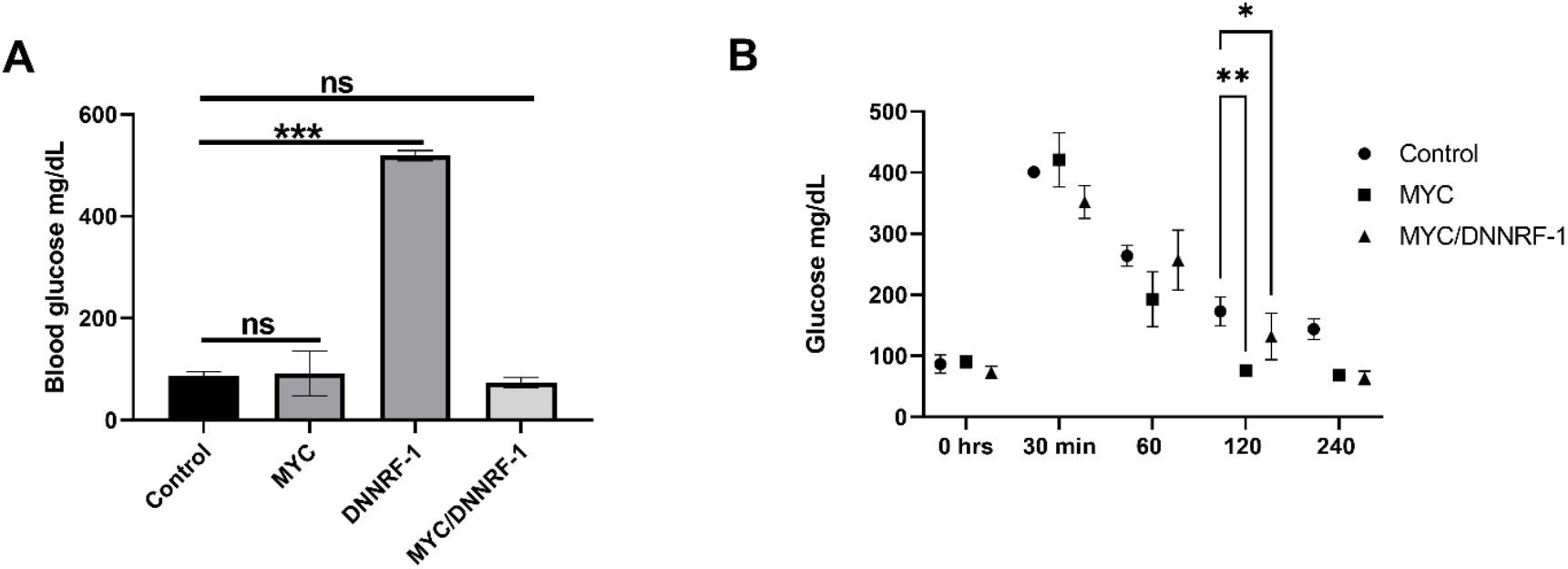
Myc corrects fasting blood glucose and glucose tolerance in pIns-DNNRF-1ER^TAM^ mice. (A) Fasting blood glucose in double transgenic pIns-*DNNRF-1ER^TAM^* /*pIns-MycER^TAM^* mice with targeted expression of both MycER and DNNRF-1ER to β cells compared with pIns-*DNNRF-1ER^TAM^* mice at 2 months of age (n=4). (B) Glucose tolerance to intraperitoneal injection of 10% glucose in PBS at a dose of 2 mg glucose/g body weight of pIns-*DNNRF-1ER^TAM^* /*pIns-MycER^TAM^* mice compared with pIns- *DNNRF-1ER^TAM^* mice at 2 months of age (n=4).

## DISCUSSION

In this study, we show that expression of a dominant-negative mutant of nuclear respiratory factor 1 (DNNRF1) in pancreatic β-cells results in the early onset of diabetes. Our data show that baseline serum insulin levels for *pIns*-*DNNRF1ER^TAM^* transgenic mice were 2-3 fold less than controls despite high levels of blood glucose. Blood glucose in *pIns*-*DNNRF1ER^TAM^* mice normalized with 1 mU/g insulin i.p., confirming that DNNRF1 expression in β cells produced insulin deficiency rather than systemic insulin resistance. The loss of insulin sensitivity found in older animals is likely a consequence of sustained hyperglycemia as observed in mouse models of type 1 diabetes and patients with T1D (Unger and Grundy, 1985; Perseghin et al., 2003; Hong et al., 2007). Although the *Ins2* promoter is known to produce expression of transgenes in hypothalamus, kidney and duodenum, reduced expression of NRF1 targets *Tfam*, *T@1m* and *T@2m* in hypothalami from DNNRF1-mice was not evident, and we could not demonstrate expression of DNNRf1 in kidney or duodenum.

Apoptotic cells were evident in transgenic islets by light and electron microscopy and immunostaining for cleaved caspase-3 confirmed an increased apoptotic rate. This differs from mice with β-cell-selective knockout of *Tfam*, a NRF1 target, which had reduced β-cell mass without evidence of apoptosis and points to a more severe phenotype when multiple NRF1 targets are inactivated (Silva et al., 2000).

In addition to apoptotic loss of β-cells, it is probable that DNNRF1 decreases glucose- stimulated insulin secretion via reduced expression of mitochondrial respiratory chain proteins regulated by NRF1 (Scarpulla, 2008). Mitochondrial dysfunction in surviving β-cells is indicated by the absence of detectable respiratory activity, low cytochrome oxidase and succinate dehydrogenase activity by histochemistry, and abnormal mitochondrial ultrastructure, without a decrease in mitochondrial content. Reduction of cytochrome oxidase and succinate dehydrogenase activities by DNNRF1 is consistent with the regulation of complex II and IV subunits by NRF1 (Dhar et al., 2008; Piantadosi and Suliman, 2008; Au and Scheffler, 1998).

Mitochondrial diabetes is associated with mtDNA mutations and is estimated to account for less than 5 percent of patients with diabetes. Despite a high penetrance of this maternally inherited form of the disease, the average age of onset is 35-40 y, raising questions about the mechanism for gradual loss of islet cell function (Maassen et al., 2004). The majority of mtDNA mutations that result in diabetes are associated with alterations in mitochondrial protein synthesis. Diabetes occurs in association with a mtDNA mutation in the tRNA-Leu gene in patients with maternally inherited diabetes and deafness (MIDD), representing 0.5-3.0% of the diabetes population. The same mutation, MTLL1*MELAS3243G, when present at higher levels of heteroplasmy (>85%) is associated with MELAS (mitochondrial encephalopathy, lactic acidosis and stroke-like episodes), which may include diabetes in its presentation (Stark and Roden, 2007). Mitochondrial myopathies associated with diabetes include mutations affecting protein synthesis at tRNA-Lys in myoclonic epilepsy and ragged red fibers (MERRF), and Kearns-Sayre syndrome and chronic progressive external ophthalmoplegia (CPEO), both deletions involving tRNAs (Austin et al., 1998). Additionally, decreased insulin secretion has been described in MIDD, Kearns-Sayre and CPEO (Piccolo et al., 1989; Becker et al., 2002; Salles et al., 2007). Surprisingly, other mtDNA mutations, rearrangements or deletions associated with inherited diseases (Leber’s hereditary optic neuropathy – ND4/6 mutation, Leigh’s syndrome/NARP (Neuropathy, Ataxia and Retinitis Pigmentosa) – ATP6 mutation) are not associated with diabetes (Maassen et al., 2004).

The selective association of mitochondrial translation mutations with diabetes suggests that NRF1 targets involved in mitochondrial translation such as *MRPS12* or mtDNA replication/ transcription (*TFAM*, *TFB1M*, *TFB2M*) may be critical for the mitochondrial diabetic phenotype (Gleyzer et al., 2005; Cam et al., 2004; Yau et al., 2021). Indeed, a common variant in *TFB1M* is associated with reduced insulin secretion, elevated postprandial glucose levels, and future risk of type 2 diabetes and analysis of *T@1m*^+/-^ mice demonstrate a reduction in mitochondrially encoded proteins (Koeck et al., 2011). In these mice, GSIS is reduced by 36%, fuel-stimulated ATP by 25% and glucose stimulated oxygen consumption by 18%. Our findings for *pIns*-*DNNRF1ER^TAM^* transgenic mice are indicative of a more severe phenotype since there is a 70% reduction in GSIS, a 60% reduction in ATP and glucose stimulated oxygen consumption was negligible. These differences are likely attributable to NRF1’s dual regulation of both mitochondrial-encoded and nuclear-encoded mitochondrial proteins and associated effects to disrupt electron transport chain protein assembly. Electron micrographs of β cells confirm that β-cells contained morphologically abnormal mitochondria filled with electron-dense matrix material obscuring the cristae. This appearance is reminiscent of mitochondria from serum-deprived cells expressing activated c-Myc that are also undergoing apoptosis and may represent misassembled proteins in the matrix prone to precipitation (Morrish et al., 2003; Margineantu et al., 2007).

We found that introduction of c-MycER^TAM^ targeted to β-cells, by generating double transgenic *pIns-DNNRF1ER^TAM^* / *pIns-c-MycER^TAM^* mice, suppressed the diabetic phenotype of pIns-DNNRF-1ER^TAM^ mice. This suggests that two pro-apoptotic factors with opposite effects on mitochondrial gene expression are capable of in vivo complementation or synergistic viability, as observed for NIH3T3 fibroblasts in vitro (Morrish et al., 2003). Direct transcription targets of c-Myc including mitochondrial genes overlapping with NRF1 targets, recently shown to require specific interactions of c-Myc with host cell factor (HCF)-1 (Morrish et al., 2008; Popay et al., 2021). The NRF1 binding motif is a non-canonical E-box that c-Myc binds *in vitro* and *in vivo*. The most likely explanation is a balancing of their effects on mitochondrial biogenesis, with restoration of mitochondrial oxidative phosphorylation and “unpriming” of mitochondrial pathways of apoptosis. The rescue of the diabetic phenotype is especially notable since both single transgenics are associated with insulin-deficient diabetes (DNNRF1 this study; c-Myc (Piantadosi and Suliman, 2008)). Recent in vitro studies have demonstrated that there is sufficient *pIns-c-MycER^TAM^* activity in the absence of tamoxifen to inhibit glucose-stimulated Ca^2+^ flux, mitochondrial hyperpolarization and insulin secretion in vitro (Pascal et al., 2008).

Although genetic polymorphisms in *NRF1* have been linked to diabetes, this study is the first study to demonstrate a direct role for NRF1 in β cell function and survival (Cho et al., 2005; Gaulton et al., 2008; Liu et al., 2008; Cam et al., 2004). This model provides a tool for future investigations into mitochondrial diabetes and opportunities to study the ameliorating effects of other regulatory factors involved in bi-genomic regulation of mitochondrial biogenesis.

### Limitations of the study

The linkage of genetic polymorphisms in *NRF1* with type 2 diabetes in Korean, Han Chinese and Finnish populations has been suggested as an example of mitochondrial diabetes. Mitochondrial dysfunction can impair glucose tolerance via two mechanisms: decreased insulin production (effects on pancreatic β-cells) or insulin resistance (effects on peripheral tissues). Mitochondrial diabetes often involves both mechanisms (Lindroos et al., 2009). Our transgenic study is limited to effects on pancreatic β-cells, and in addition to potential differences in the degree of NRF1 functional deficits between the mouse model and germline polymorphisms, is not designed to replicate the genetic association in human disease.

Analysis of pancreatic islet composition and glucose-stimulated insulin secretion support a role for DNNRF1-dependent mitochondrial dysfunction in reduced insulin secretion in viable β- cells and in progressive loss of β-cells via apoptosis. The relative contribution of these mechanisms may depend on the extent of mitochondrial dysfunction, and our study did not attempt to explore additional mouse models with more limited NRF1 impairment. Limited studies of islets obtained at autopsy in mitochondrial diabetes have also demonstrated β-cell depletion (Kobayashi et al., 1997).

We proposed one mechanism of mitochondrial dysfunction to be misassembly of mitochondrial electron transport chain complexes. While the appearance of diffusely electron- dense material in the mitochondrial matrix has been associated with reduced detergent extractability (Margineantu et al., 2007), we did not specifically test for the presence of incomplete ETC complexes in β-cells from DNNRF1-mice.

## ACKNOWLEDGMENTS

We thank Hockenbery and Sweet lab members for helpful discussions and Bonnie Kraskouskas for advice and technical assistance. Support was provided by NIH CA158921 (D.M.H.), DK17047, DK063986 (I.R.S.) and National Science Foundation (IIP-0750508) (I.R.S.).

## AUTHOR CONTRIBUTIONS

F.M. and D.M.H. conceived the project. F.M., H.G., J.N., and L.H. performed all experiments on live mice, I.R.S. and I.T.K. isolated islets and performed islet perfusion studies, and S.E.K. advised on immunohistochemistry pathology markers and evaluated results of immunohistochemistry assays.

## Resource availability

### Lead Contact

Dr. David Hockenbery

## Materials Availability Statement

Transgenic DNNRF1 mice are available as cryogenic sperm and plasmids are available upon request.

## Data and code availability

- All analyzed and raw data reported in this paper will be shared by the lead contact upon request.
- This paper does not report original code.
- Any additional information required to reanalyze the data reported in this paper is available from the lead contact upon request.

## Experimental model and study participant details

C57BL/6J is the most widely used inbred strain for transgenic research and were used at age 6-8 weeks. The mice were kept at the animal facility of the Fred Hutchinson Cancer Center with 12/12h light/dark cycles at 65-75^0^F with 40-60% humidity in approved housing in accordance with the guidelines of the FHCRC Institutional Animal Care and Use Committee (File: 1238).

## Method Details

### Plasmid construction

A truncated NRF-1 with the transcriptional activation site removed and fused to the hormone-binding domain of a modified estrogen receptor, DNNRF-1ERTAM, has been described previously (Morrish et al., 2003; Wu et al., 1999; Gonen and Assaraf, 2010; Tokusumi et al., 2004; 12 Kherrouche et al., 2004). The rat insulin promoter expression plasmid was a gift from G. Evan (UCSF) and comprises 0.7 kb of the rat insulin promoter and 1.6 kb of the β-globin polyadenylation sequence in a pUC18 backbone (pIns) (Ohashi et al., 1991). DNNRF-1ERTAM DNA was excised from pBabe vector by partial BamH1/BstI digest and blunt-ended with T4 DNA polymerase. The pIns plasmid was cut with BamHI and blunt-ended with T4 polymerase, and ligated to the DNNRF-1ERTAM sequence. Since no unique restriction sites were present in the pUC18 vector, the insert was excised with SmaI and HindIII to generate two fragments (SmaISmaI and SmaI-HindIII) that were subsequently sub-cloned into the SmaI-HindIII site of pBluescript in a 3-way ligation. The final transgene construct was then excised from pBluescript as a 4.25 kb NotI-HindIII fragment. Purified DNA was resuspended for pronuclear injection in sterile buffer (10 mM Tris, 0.1 mM EDTA pH 7.4) at a concentration of 5 ng/μl.

### Generation of pIns-DNNRF-1ER TAM transgenic mice

pIns-DNNRF-1ERTAM DNA was injected into male pronuclei of day 1-fertilized (CBA × C57BL/6) F1 embryos. Injected embryos were transferred into day 1-plugged pseudopregnant foster mice, and the litters were screened for presence of the transgene by PCR. Founder mice were backcrossed to C57BL/6 for > 5 generations to establish transgenic lines. For Southern blot analysis, genomic DNA isolated from mouse tails was digested overnight with BamHI or EcoR1, fractionated on a 1% agarose gel, transferred to a Hybond-N+ filter (Amersham), and probed with a 32P-labeled 0.5 kb human NRF-1 cDNA fragment. Transgene copy number was estimated by densitometry with dilutions of a NotI/HindIII transgene fragment. All subsequent genotyping was done by PCR using the following primer sequences that span DNNRF-1 and the ERTAM (forward 5’ATCAGCAAACGCAAACAC3’ and reverse 5’GCCAGACGAGACCAATCAT C3’). Littermates, or age-matched C57BL/6 mice were used as controls in all experiments.

### Generation of pIns-DNNRF-1ER TAM/ pIns-MycER TAM double transgenic mice

pIns-MycER TAM mice (kindly provided by Gerard Evan, UCSF) were crossed with pInsDNNRF-1ER TAM mice and screened for the presence of both transgenes using a primer set 13 consisting of a common 3’ primer for the ER region (5’TAGATCATGGGCGGTTCAGC3’) and specific 5’ primers for DNNRF-1 (as above) and Myc (5’CAAGAGGGTCAAGTT GGACA3’). These primer sets amplified PCR products of 683 bp and 422 bp, respectively.

### Histological, immunohistochemical and histochemical analysis of pancreatic tissue

Pancreata were excised from mice, and 5 mm pieces of tissue were fixed overnight in 10% neutral-buffered formalin, embedded in paraffin wax, and sectioned (5–10 μm).

Immunohistochemical staining and morphometric analysis were performed in tissue sections separated by 200 μM to include > 20 islets. Apoptotic cells were identified in paraffin-embedded sections by immuno-staining with activated cleaved caspase 3 antibody (Cell Signaling Technologies, Inc., Danvers, MA). Immunohistochemistry was performed using guinea pig antiinsulin primary antibody (Fitzgerald Labs, Concord, MA, USA) diluted 1:450, rabbit anti-glucagon primary antibody (Vector Labs, Burlingame, CA) diluted 1:30, visinin-like protein 1 primary antibody (Thermo Scientific Pierce Products) diluted 1:100. Immunostaining was completed using a three-step streptavidin technique, as described (Carson and Hadik, 2004).

Histochemistry for cytochrome oxidase and succinate dehydrogenase activities were performed on formalinfixed and cryosections, respectively, using methods previously described (Morrish et al., 2003; Sweet et al., 2004).

### Perifusion system for islets

Islets were harvested and a flow chamber was used to evaluate dynamic oxygen consumption and insulin release by islets in response to glucose, as previously described (Niswender et al., 2005).

### Static measurement of insulin secretion

Static insulin secretion was determined with different secretagogues as previously described (Patti et al., 2003). Briefly, islets were transferred into a Petri dish containing Krebs Ringer buffer (KRB), 0.1% BSA and 3 mM glucose and incubated at 37°C in 5% CO2 for 60 min. 14 Subsequently, islets were placed into 96-well plates (10 islets/well) containing no glucose, 20 mM glucose, 30 mM KCl, or 20 mM glucose and 30 mM KCl, and incubated for 60 min. At the end of this period, supernatant was assayed for insulin using an RIA kit (Linco Research Inc.). For solutions containing 30 mM KCl, the NaCl concentration in the KRB was reduced by 25 mM.

### Electron microscopy

Islets were isolated and fixed for electron microscopy as previously described (Morrish et al., 2003). Cell and mitochondrial areas were calculated using the NIH image processing program ImageJ. Image J measurements Images were imported into Image J and the area of each islet was measured using the freehand selection tool. The area in Image J is provided as square pixels with an aspect ratio of 1. For analysis of the cell density in an islet, nuclei were counted and divided by the total area of an islet and the data expressed as nuclei per square pixel. Caspase and insulin immunostaining was quantitated as a percentage of islet area. Mitochondria and insulin secretory granules were quantitated per unit cellular area.

### Animals

This study was carried out in strict compliance with the recommendations in the Guide for the Care and Use of Laboratory Animals of the National Institutes of Health. All mouse protocols used in this study were approved by the FHCRC Institutional Animal Care and Use Committee (File #1238). Mice were housed at the FHCRC and fed a regular chow diet. All surgery was performed under sodium pentobarbital anesthesia and all efforts were made to minimize suffering.

### Clinical chemistry assays

Blood or urine glucose levels were measured with a portable glucose measuring device (OneTouch, LifeScan, Milpitas, CA), accurate for glucose values between 10 to 600 mg/dl. Serum insulin levels were measured by ELISA (Mouse Phenotyping Center, University of Washington, WA). All animals used in these studies were between 2 and 8 months of age. For fasting blood glucose measurements, mice were fasted for 4 h and then placed in a heated chamber (32°C) for 10 min to induce vasodilation. Blood (0.2 ml) was collected from tail veins. ATP assays ATP content in freshly isolated islets was measured by the CellTiter-Glo assay (Promega).

### Glucose tolerance test and insulin sensitivity assays

For the glucose tolerance test (GTT), mice were fasted overnight (14 h) and injected intraperitoneally with 10% glucose in PBS at a dose of 2 mg glucose/g body wt. For the insulin sensitivity assay, mice were fasted overnight and injected intraperitoneally with 0.1 units/ml Humulin R insulin (Lilly, Indianapolis, IN) in sterile PBS at a dose of 1.0 mU insulin/g body wt.

### Quantification and statistical analysis

Two-tailed ANOVA with correction for multiple comparisons and Student’s T-tests were conducted in Prism version 9, and the statistical significance, and values of n are indicated in the figure legends. Data in the figures are represented of 2-3 replicate experiments and expressed as mean +/- SEM and significance is marked with *=p<0.05, **=p<0.01, ***=p<0.001. A minimum of 10 images were used for IHC quantitation and we acknowledge potential bias in these data as the researcher was not blinded.

### Graphic illustrations

The graphic illustrations displayed in this paper were created with Prism and Biorender.

