## Supplemental files for "Mitochondrial diabetes in mice expressing a dominant-negative allele of nuclear respiratory factor-1 (*Nrf1*) in pancreatic β-cells"

***Supplemental Table and Figure legend***

**Supplemental Table 1.** 2^ΔCT values from qRT-PCR analysis of NRF-1 target genes in islets and hypothalamus. Results for qRT-PCR analysis. Data are mean and SE values derived from analysis of up to 3 independent wild type or DNNRF-1 mice. Mice were age and sex matched.

**Supplemental Table 2.** Summary of primers used for qRT-PCR.

***Supplemental figure legend***

**Supplemental Figure 1**. *Onset of increase in non-fasting blood glucose in DNNRF-1 mice and islet morphology.*

Blood glucose levels measured in control and DNNRF-1 mice at 3, 5 and 8 weeks of age (n=8) (A). Quantitative analysis of (B) islet area, (C) nuclei per unit islet area, (D) active caspase-3 immunostaining as percent of islet area and (E) insulin immunostaining as percent of islet area in 2 month (black bar) and 6 month-old (white bar) mice . Data from n= 3 mice per group with analysis of 20 islets per pancreas from sections separated by 200 μM.

**Supplemental Figure 2.** *Insulin and glucagon immunohistochemistry.*

Insulin (A) and glucagon (B) immunostaining for control mice. Insulin (C) and glucagon (D) staining for pIns-DNNRF-1ERTAM mice (n=3).

**Supplemental Figure 3.** *Insulin and visinin-like 1 protein immunohistochemistry.*

Insulin (A) and visinin-like 1 (B) immunostaining for control mice. Insulin (C) and visinin-like 1 (D) staining for pIns-*DNNRF-1ER$^{TAM}$* mice (n=3).

**Supplemental Figure 4.** *Morphometric analysis of β-cell electron micrographs.*

(A) Cellular area of pIns-*DNNRF-1ER$^{TAM}$* and wildtype β-cells. (B) Mitochondria per cell. (C) Mitochondrial area normalized to cell area. (D) Insulin secretory granules normalized to cell area. (n=10).

**Supplemental Figure 5.** *Static islet assay for response to glucose and KCl.*

Each static analysis experiment was performed with four replicates of 10 islets per well, treated with glucose at 0 mM (black bar), 20 mM (white bar), 0 mM plus 30 mM KCl (gray bar) and 20 mM glucose plus 30 mM KCl (cross-hatched bar). The table included in this figure provides p-values for t-test comparison and ANOVA with multiple comparison data for control (wild-type) versus DNNRF-1 mice.

**Supplemental Figure 6.** *ATP levels in isolated islets.*

ATP content in freshly isolated islets was measured by the CellTiter-Glo assay. Data are means of 3 replicate experiments. For each experiment, ATP was measured from 3 islets in triplicate (controls) and duplicate (transgenics).

**Supplemental Figure 7***. Islet morphology in pIns-MycER$^{TAM}$/pIns-DNNRF-1ER$^{TAM}$ mice.*

H&E staining of pancreatic islet from non-transgenic control (A) and pIns-*DNNRF-1ER$^{TAM}$* /pIns-*MycER$^{TAM}$* transgenic mice (B) at 2 months of age. Insulin staining of pancreatic islets from control (C) and pIns-*DNNRF-1ER$^{TAM}$* /pIns-*MycER$^{TAM}$* (D) mice at 2 months of age. 40 X magnification (n=3).

**Supplemental Figure 8.** *Genotyping of Myc/DNNRF-1 mice.*

An example of genotyping for a litter of mice generated from a cross between Myc and DNNRF-1 mice. Mice positive for both genes generate two products representing the presence of the Myc transgene (683 bp) and the DNNRF-1 transgene (422 bp). Wildtype mice do not produce any product and single transgenics produce a single PRC product.

**Supplemental Figure 9.** *Mitochondrial number and area.*

(A) Data for the number of mitochondria per cell derived from electron micrographs (n=5). (B) Data for the sum of the mitochondrial area/cell area (n=5).

Supplemental Table 1

| Islet | | | Hypothalamus | | |
|---|---|---|---|---|---|
| | DNNRF-1 | p-value | | DNNRF-1 | p-value |
| *Tfam* | 0.19+/-0.09 | 0.00004 | *Tfam* | 1.08+/-0.43 | 0.38 |
| *Tfb1m* | 0.16+/-0.11 | 0.0001 | *Tfb1m* | 0.99+/-0.45 | 0.49 |
| *Tfb2m* | 0.16+/-0.12 | 0.00007 | *Tfb2m* | 1.07+/-0.29 | 0.35 |

Supplemental Table 2

| Gene | Accession number | Reference | Product size (bp) | Forward primer | Reverse primer |
|---|---|---|---|---|---|
| *TBP* | NM_013684 | PMID:9891007 | 190 | ACCCTTCACCAATGACTCCTATG | ATGATGACTGCAGCAAATCGC |
| *Tfam* | NM_009360 | PMID:20107496 | 175 | CAAGTCAGCTGATGGGTATGG | TTTCCCTGAGCCGAATCATCC |
| *Tfb1m* | BC032930 | PMID:20107496 | 189 | AATTTCCTCCTGGACTTGAGG | AGAGAGCATCTGTAACCCTGG |
| *Tfb2m* | NM_008249 | PMID:20107496 | 201 | GTTCGAATGACTCCTCGTAGG | CATTCTAGCAGCTGTGTCTCC |
| *DNNRF-1* | Fusion protein | This study | 100 | CAGCCACACATAGTATAGCTCATCTTG | GAAGTCCTCTGTACTCCCGACG |
| *Cox5b* | NM_009942 | PMID: 18222924 | 217 | CAAGGTTACTTCGCGGAGTG | TCCTTGGTGCCTGAAGCTG |
| *Cycs* | NM_007808 | PMID: 18222924 | 177 | TTCAGAAGTGTGCCCAGTGC | CTCCAAATACTCCATCAGGGTATC |

Supplemental Figure 1

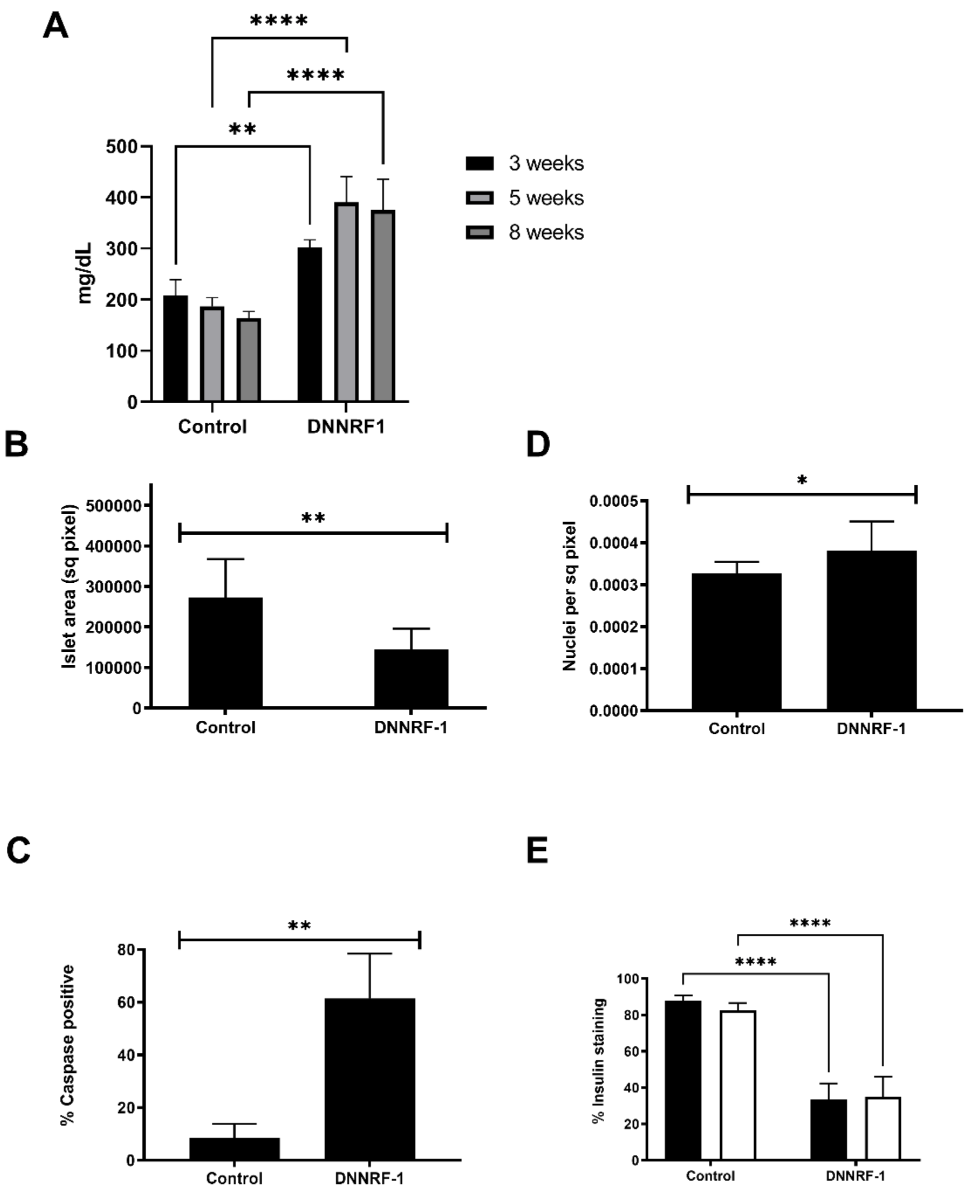

A
****
****
**
500
400
300
200
100
0
mg/dL
Control
DNNRF1
3 weeks
5 weeks
8 weeks
B
**
500000
400000
300000
200000
100000
0
Islet area (sq pixel)
Control
DNNRF-1
D
*
0.0005
0.0004
0.0003
0.0002
0.0001
0.0000
Nuclei per sq pixel
Control
DNNRF-1
C
**
80
60
40
20
0
% Caspase positive
Control
DNNRF-1
E
****
****
100
80
60
40
20
0
% Insulin staining
Control
DNNRF-1

Supplemental Figure 2

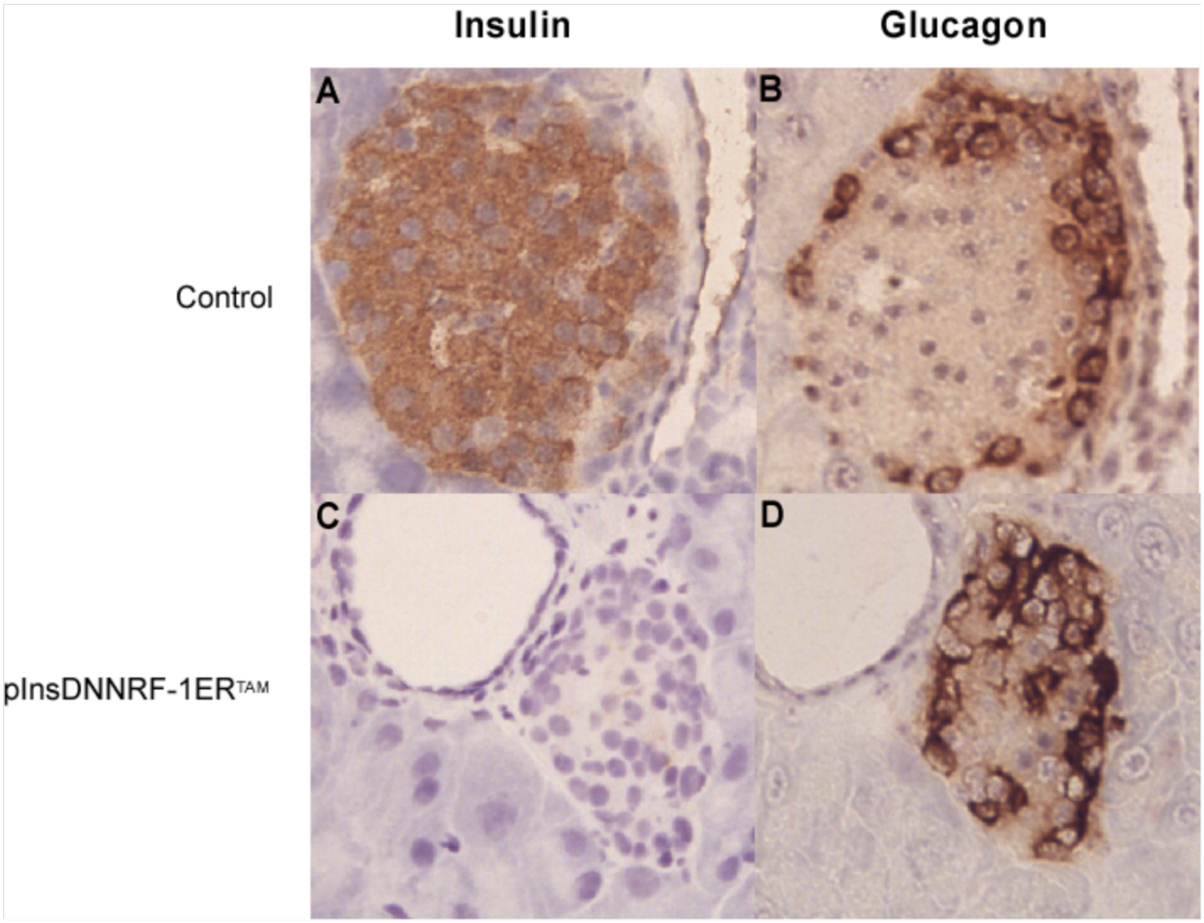

Supplemental Figure 3

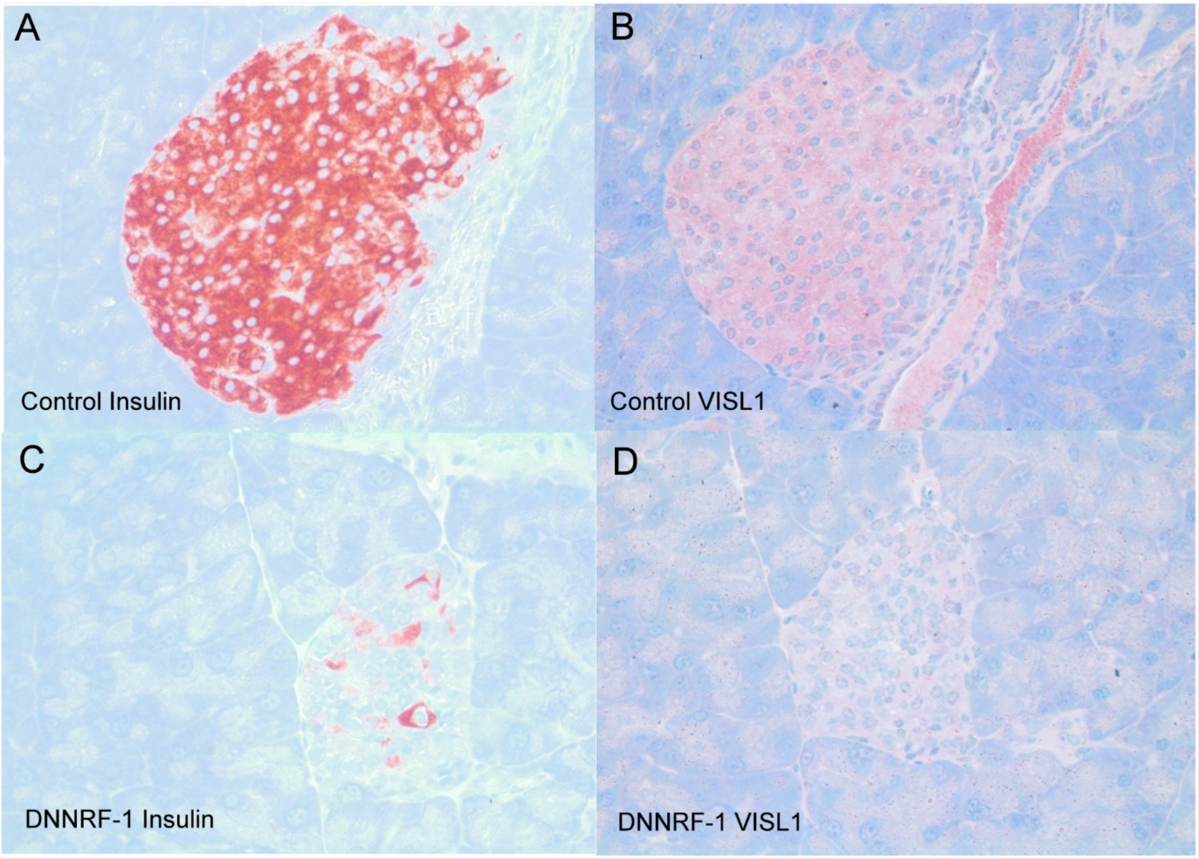

Supplemental Figure 4

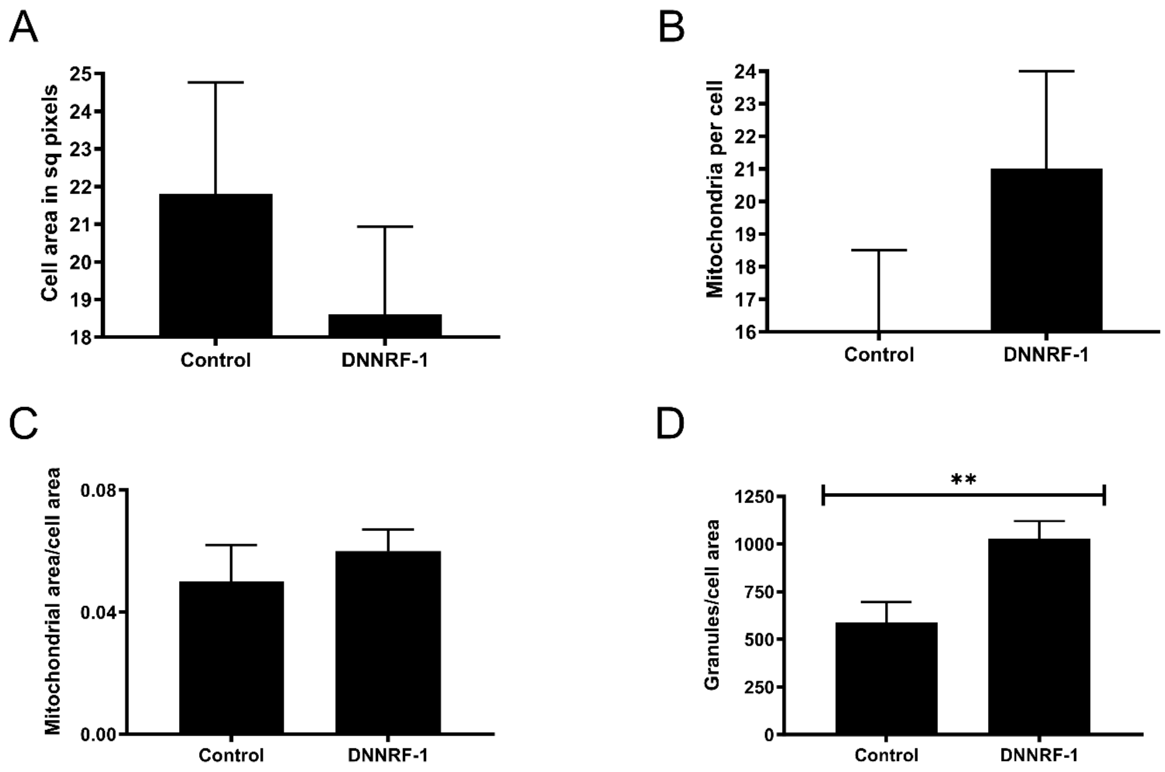


Supplemental Figure 5

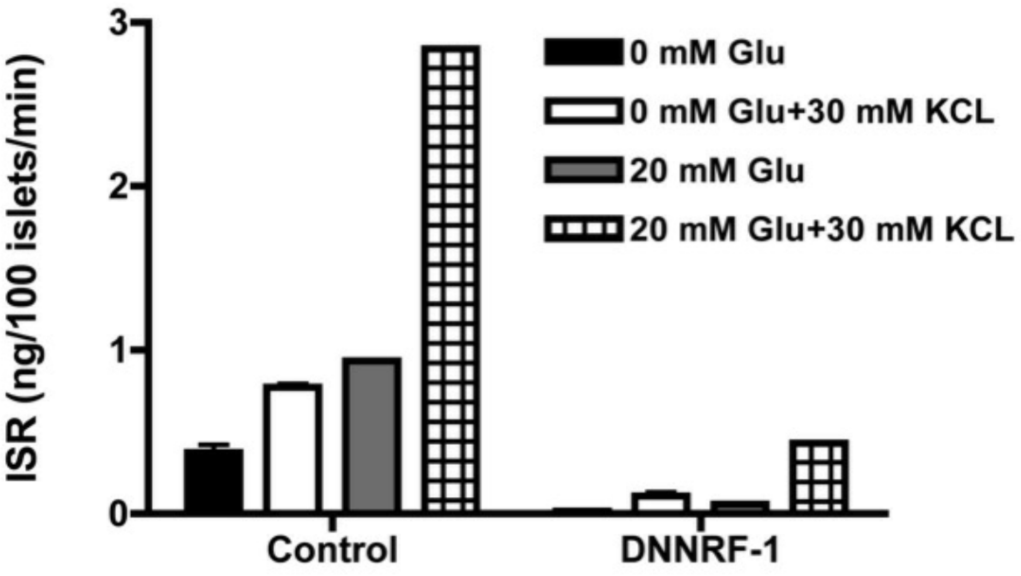


| t-test Wt vs DNNRF-1 | |
|---|---|
| | t-test p values |
| 0 mM Glu | 0.01 |
| 0mM Glu+ 30 mM KCL | <0.0001 |
| 20 mM Glu | <0.0001 |
| 20 mM Glu+30 mM KCL | 0.005 |

| ANOVA | Wild-type | DNNRF-1 |
|---|---|---|
| 0 mM Glu vs 0mM Glu+ 30 mM KCL | P > 0.05 | P > 0.05 |
| 0 mM Glu vs 20 mM Glu | P > 0.05 | P > 0.05 |
| 0 mM Glu vs 20 mM Glu+30 mM KCL | P < 0.001 | P < 0.01 |
| 0mM Glu+ 30 mM KCL vs 20 mM Glu | P > 0.05 | P > 0.05 |
| 0mM Glu+ 30 mM KCL vs 20 mM Glu+30 mM KCL | P < 0.01 | P < 0.05 |
| 20 mM Glu vs 20 mM Glu+30 mM KCL | P < 0.01 | P < 0.01 |

Supplemental Figure 6

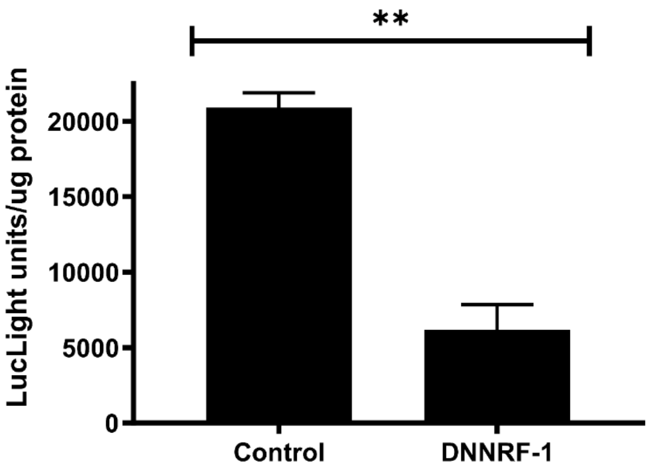


Supplemental Figure 7

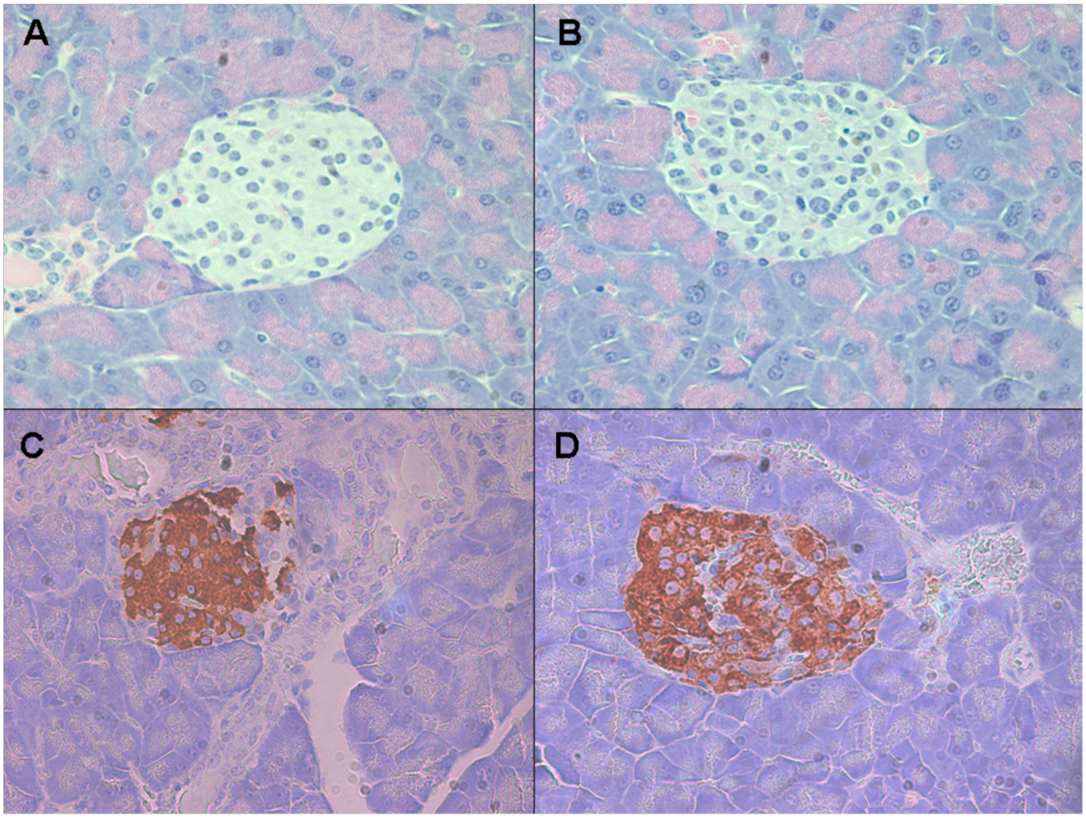
A
B
C
D

Supplemental Figure 8

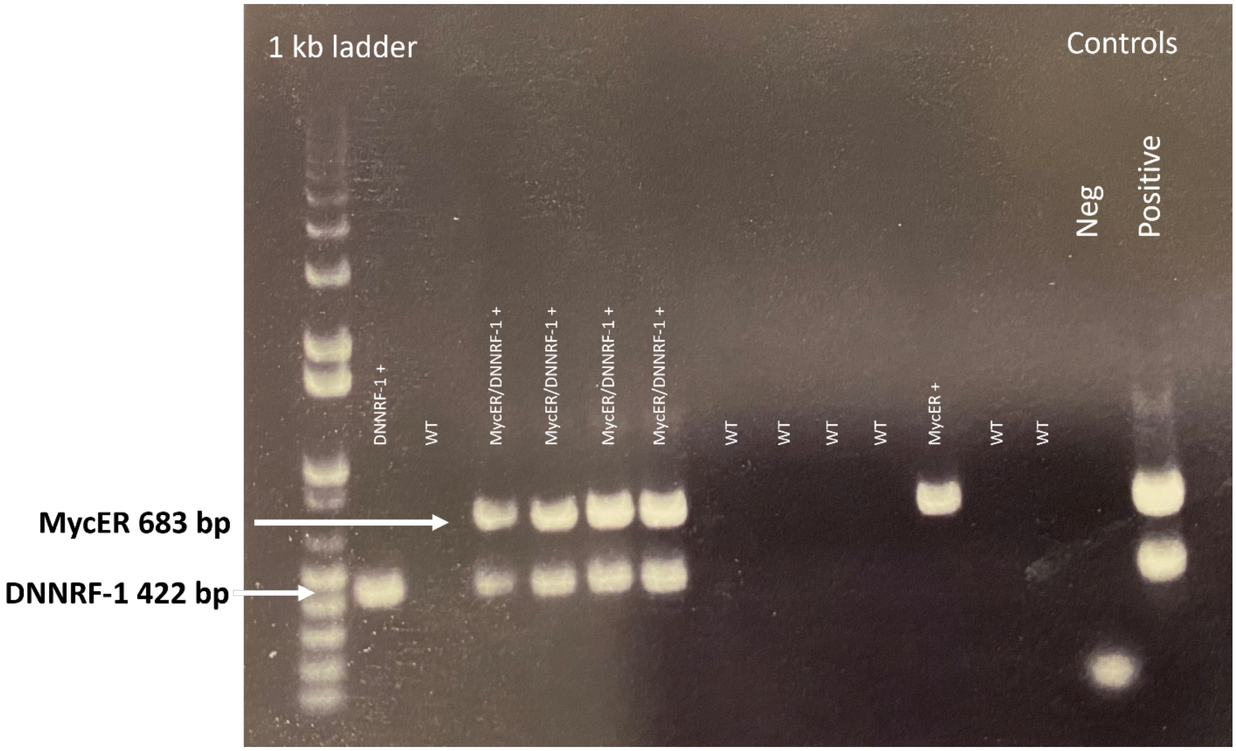

Supplemental Figure 9

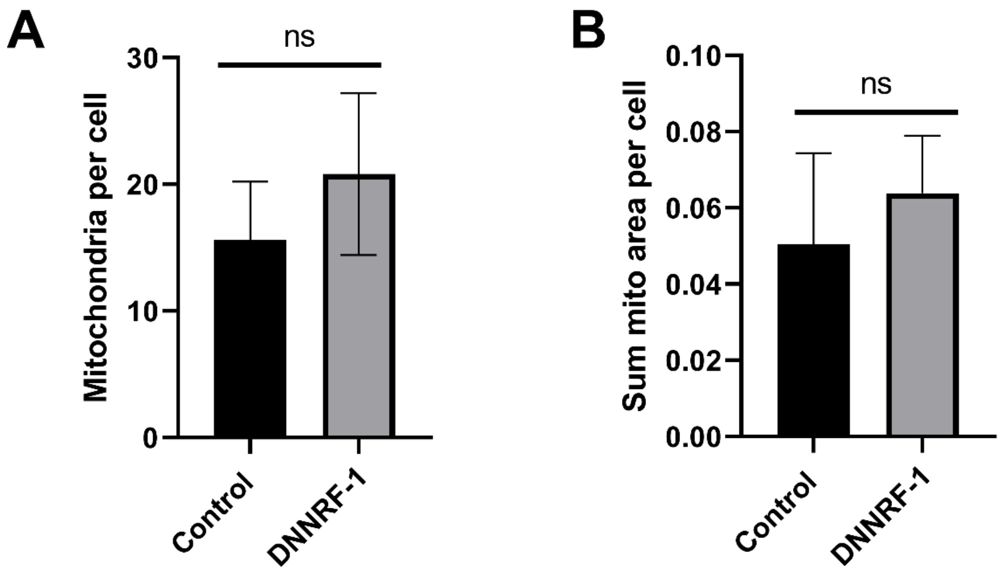
